# Computational Roles of Intrinsic Synaptic Dynamics

**DOI:** 10.1101/2021.04.22.441034

**Authors:** Genki Shimizu, Kensuke Yoshida, Haruo Kasai, Taro Toyoizumi

## Abstract

Conventional theories assume that long-term information storage in the brain is implemented by modifying synaptic efficacy. Recent experimental findings challenge this view by demonstrating that dendritic spine sizes, or their corresponding synaptic weights, are highly volatile even in the absence of neural activity. Here we review previous computational works on the roles of these intrinsic synaptic dynamics. We first present the possibility for neuronal networks to sustain stable performance in their presence and we then hypothesize that intrinsic dynamics could be more than mere noise to withstand, but they may improve information processing in the brain.

**Highlights:**

- Synapses exhibit changes due to intrinsic as well as extrinsic dynamics
- Computational frameworks suggest stable network performance despite intrinsic changes
- Intrinsic dynamics might be beneficial to information processing

## Introduction

Learning is a major area of research in both neuroscience and machine learning. Synaptic plasticity, changes in synaptic weight depending on the activity of neurons, is an essential mechanism for learning and memory in the brain [1]. One of the prevailing properties of synaptic plasticity is characterized by Hebb’s rule [2,3], in which synaptic weights change according to the correlation between pre- and postsynaptic neuronal activities. Hebbian plasticity, including long-term potentiation and depression [1] and variants such as spike-timing-dependent plasticity (STDP) [4,5], has been proposed to contribute to efficient information processing in the brain. Further, being originally inspired by operations of neurons in the brain, artificial neural networks (ANNs) have been investigated as a promising subject in machine learning. Synaptic weights are trained in ANNs by an algorithm to achieve optimal performance (e.g., by the error back-propagation rule [6]). Although there are differences in the learning rules used in the brain and ANNs [7], the importance of changing the synaptic weight is established.

The postsynaptic counterpart of the synapse that receives a signal from the presynaptic neuron is called the dendritic spine, a small protrusion from a neuron’s dendritic shaft (Figure 1A) [8,9]. Previous experimental results showed that the spine volume of a synapse is closely correlated with its weight [10–13], and synaptic plasticity protocols such as long-term potentiation and depression [14–17] and STDP [18,19] induce spine enlargement or shrinkage. Several in vivo studies have demonstrated that behavioral learning involves change in spine volumes and numbers [20–22] and that the selective shrinkage of the spines potentiated during training causes memory erasure [23]. These experiments give strong evidence that spines play a fundamental role in both learning and memory via Hebbian plasticity.

**Figure 1.**
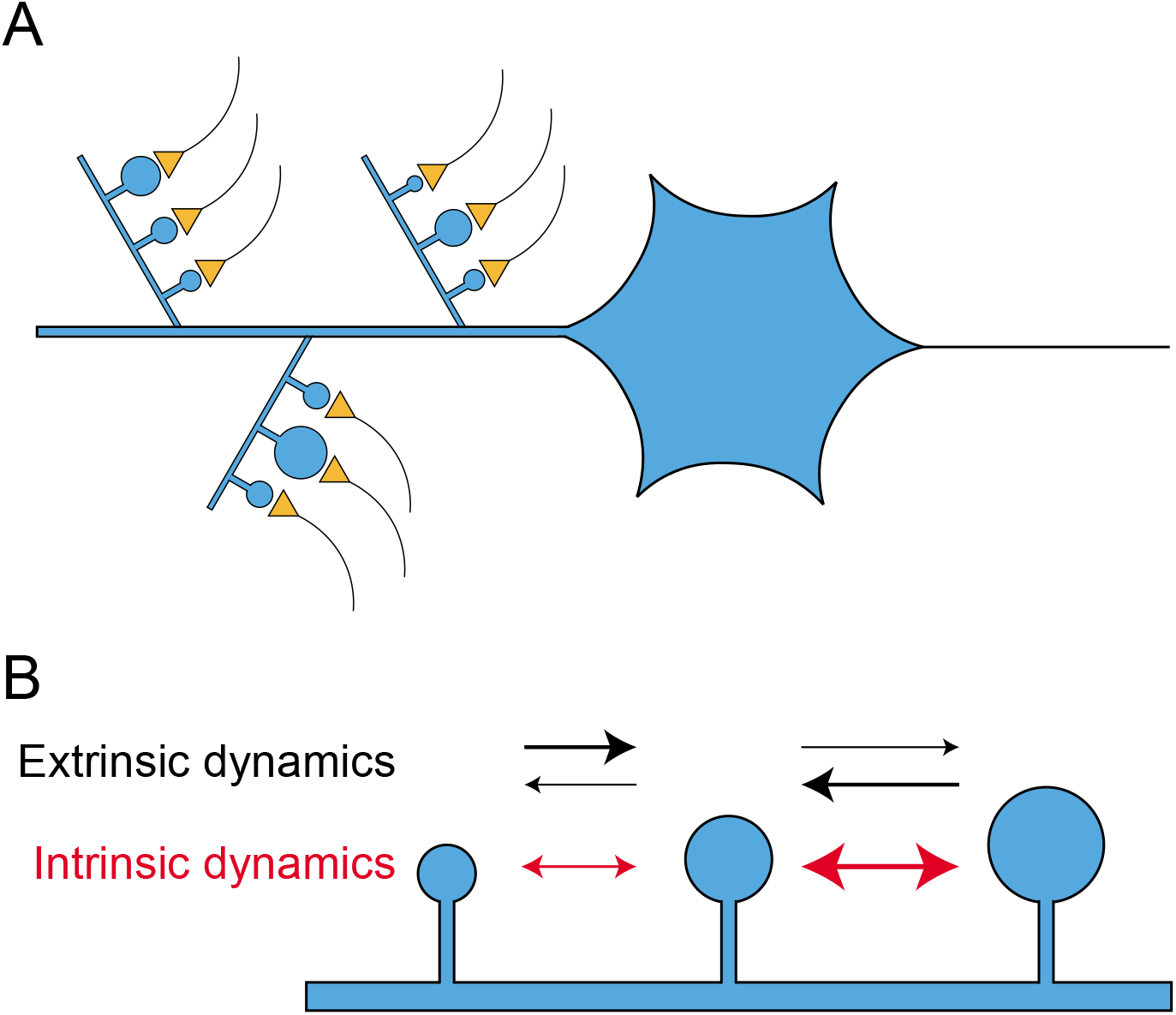
A schematic image of dendritic spines and spine fluctuations. A. Presynaptic neurons transmit information to postsynaptic neurons through synapses. The postsynaptic sites of excitatory synapses mostly form dendritic spines, small protrusions on dendritic shafts. Synaptic weights are nearly in proportion to spine volumes, and potentiation/depression of synapses are associated with enlargement/shrinkage of dendritic spines. B. The spine size is regulated by two factors: extrinsic dynamics (activity-dependent plasticity) that reflects neuronal activities and intrinsic dynamics (fluctuation) that is independent of neuronal activities. Note that for larger spines intrinsic dynamics are dominant, while smaller spines are rather subject to extrinsic dynamics.

However, many studies have reported that spines are highly volatile (Figure 1B) [24,25]. About 40% of the spines in the hippocampus are replaced within 4 days [26] and more than 50% of the spines in the auditory cortex are replaced within two weeks [27]. The volume of existing spines also exhibits substantial variability [28]. Although this variability can partly be explained by the effect of neuronal activities, spine volumes change significantly in an activity-independent manner as well. Several studies reported that spine sizes [29] or postsynaptic density molecules [30] fluctuate even under the condition where neuronal activities were inhibited pharmacologically. Spine formation and elimination under the suppression of neuronal activities were also observed in *in vivo* studies [31]. Surprisingly, one study reported that more than half of the changes of individual spines are estimated to occur independently of the neuronal activities by analyzing the size correlation of spines between the same pre- and postsynaptic neurons [32]. Note that, in contrast to the intrinsic dynamics, we call synaptic changes due to activity-dependent synaptic plasticity extrinsic synaptic dynamics hereafter.

The existence of intrinsic synaptic dynamics raises two important questions. First, how is memory retained in the presence of intrinsic dynamics? While many studies have attributed memory to synaptic weight, some recent works have suggested stability of memory under spine fluctuations. Second, do intrinsic dynamics provide benefits for information processing in the brain? Generally, fluctuations can play an important role in biological processing such as selecting a suitable gene expression attractor with noise [33] and allowing molecular motors to transport cargoes robustly exploiting thermal Brownian components [34]. In the brain, recent theoretical and experimental studies on intrinsic spine dynamics further suggest their benefits for statistical inference, stable memory, and exploration in reinforcement learning. We address these questions by reviewing several studies on intrinsic spine dynamics.

## Memory could be stable despite synaptic fluctuations

Previous studies have proposed different theoretical perspectives on how memory could be stable despite intrinsic synaptic dynamics. We begin this section by discussing the possibility that memory might be stored in synapses that are little affected by intrinsic dynamics. We then focus on the paradigms in which neural computation primarily relies on the network architecture rather than individual synaptic weights. Finally, we discuss that the effect of intrinsic dynamics could partially be canceled by extrinsic synaptic dynamics. These perspectives are not mutually exclusive but can cooperate towards stability in the presence of volatility.

### Intrinsic dynamics might be harmless

If intrinsic dynamics do not affect synapses that essentially store memory, the stability of memory is not surprising. For example, dendritic spines are more stable in the cortex than in the hippocampus [26,27]; within the same area, the lifetime of spines becomes longer with their size, age, or morphological and functional maturation [27,35]. Thus long-term memory might be maintained via those relatively long-lasting synapses. Computational models which predict the stability of only a subset of synapses for memory storage support this idea [36–38]. For example, Mongillo et al. [38] showed that rewiring of excitatory connections had much less influence on memory compared with rewiring of inhibitory connections in the balanced spiking neural network model of the cortical circuit (Figure 2A). Their result arises from the condition that, under the balanced regime, the firing rates of the inhibitory neurons are higher and more diverse than the excitatory counterpart and thus an inhibitory synapse has a greater influence on neural activity than an excitatory synapse on average.

**Figure 2.**
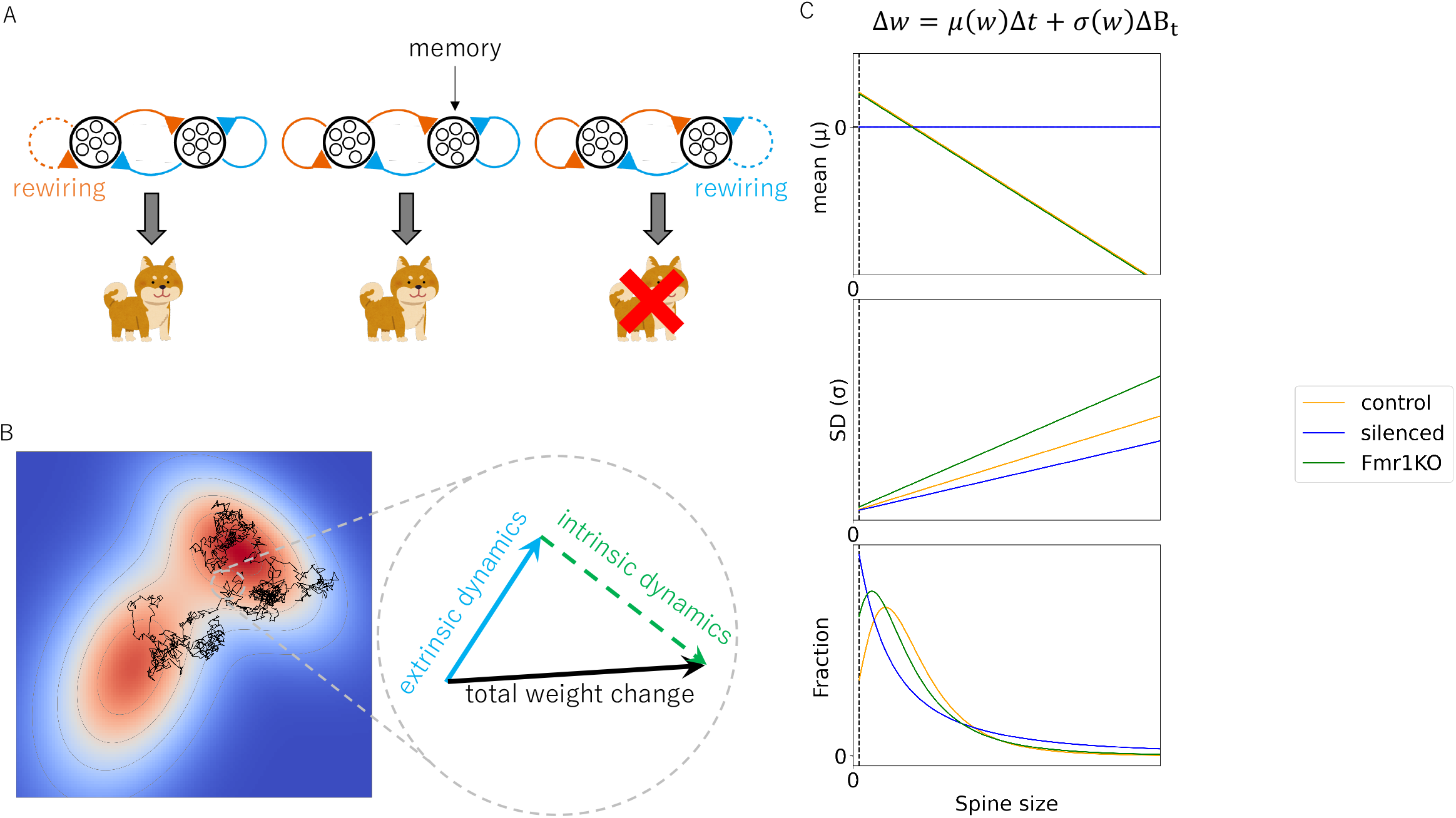
Examples of various computational perspectives on intrinsic dynamics. A. Information might be mainly stored in the subset of neurons and synapses and modifying the rest could be harmless. In the schema, memory is stored in the neurons and synapses in the right population (blue). As a result, rewiring synaptic connections in the right population impairs memory, while rewiring in the left (orange) does not. For example, in the case of computational modeling work by [38], rewiring excitatory synapses did not hinder memory recall whereas rewiring inhibitory synapses did. Note that they used a balanced cortical network model with the average firing rate higher and more diverse in inhibitory neurons, and also they assumed that cell assemblies were formed via anti-Hebbian plasticity of inhibitory synapses as well as Hebbian plasticity of excitatory neurons. B. It is widely accepted that the brain represents and manipulates uncertainty in various ways. The framework proposed by Kappel et al. [66,67] suggests that a combination of extrinsic and intrinsic dynamics could work as sampling dynamics from the target distribution of possible network connectivity. C. The skewed and heavy-tailed distribution of synaptic weights naturally arises from modeling synaptic dynamics as a stochastic process. If we use Brownian motion to describe the randomness of synaptic dynamics, fluctuations with an average change *μ*(*w*) and a standard deviation *σ*(*w*) can be described by a stochastic differential equation *dw* = *μ*(*w*)*dt* + *σ*(*w*)*dB_t_* (where *B_t_* represents standard Brownian motion). The drifting term *μ*(*w*) and diffusion term *σ*(*w*) were measured in various conditions, with which the stationary distributions of synaptic weights could be obtained. (orange): control condition [29,86]. (blue): silenced condition [29]. (green): Fmr1 knockout (KO) condition, which is a mouse model for autism spectrum disorder (ASD) [86]. The distributions have a common skewed and heavy-tailed shape in all conditions. The smaller spines observed in the Fmr1 KO model are explained by the enhanced spine turnover (larger *σ*(*w*)). Note that the plots are schematic illustrations that approximate but do not exactly replicate original fittings.

Another line of works focuses on the prevalence of multisynaptic connection [39]. While traces of extrinsic dynamics naturally correlate among synapses between the same pair of neurons, intrinsic dynamics are independent of each other. Thus creating multiple synaptic contacts could partly cancel the perturbation caused by intrinsic dynamics [40–42].

### Is it connectome that matters?

Another perspective emphasizes the importance of finding a suitable structure of network connectivity rather than optimizing each synaptic weight parameter. If intrinsic dynamics preserve this structure, fluctuation in weights could be insignificant. In the field of neuroscience, connectome studies aim to draw a comprehensive “wiring diagram” of the brain [43]. It has been reported that various features of network architectures, both local and global, might be well preserved among individuals or species [44] and that abnormal connectivity is linked to psychiatric disorders [45]. This reproducibility of the connectome hints that the network structure might play a major role in neural computation, separately from individual synaptic weights.

In accordance with this idea, recent works have shown in various settings that neural networks with appropriately designated architectures for specific tasks can perform reasonably even with random initialization. For example, two well-known building blocks in ANNs, convolutional neural network (CNN) (for visual image [46]) and long short-term memory (LSTM) (for processing time-series data [47]), are capable of effective image processing [48,49] and time series prediction [50], respectively, with random assignment of connectivity. Inspired by these results, Gaier and Ha [51] proposed a generic algorithm to search for architectures that could perform well over a wide range of weights. Their algorithm is a variant of evolutionary computing for topology search, starting from simple initial status and evolving network architectures incrementally based on their average performance over a set of different weight parameters. Another example where individual synaptic weights are insignificant for neural computation is found in [52]. This work demonstrated that low-rank structures in recurrent networks can be essential for task-performance, where resampling the network connectivity while keeping the underlying structures led to a comparable performance. Altogether, these works raise the possibility that fluctuations in synaptic weights can be mitigated as long as the qualitative characteristic of network connectivity is preserved.

### Intrinsic dynamics might be continually corrected by extrinsic dynamics

We conclude this section by mentioning that memory could be maintained via continual correction by extrinsic dynamics [53–55]. This possibility has long been examined in theoretical settings (e.g. [56]) and recent modeling works inspect it in more true-to-life situations, showing that cell assembly (an ensemble of neurons whose coherent activity represents memory) [54] or neural representation [55] is maintained if activity-dependent plasticity counteracts synaptic changes induced by intrinsic dynamics. While the theories in the previous sections assume that a part of neural circuit components evades intrinsic dynamics, this paradigm can apply to the situation where intrinsic dynamics affect every synapse.

## Intrinsic dynamics could contribute to brain function

In the previous section, we reviewed work supporting the idea that (some) synaptic fluctuations may be harmless under certain conditions. An implicit assumption behind them is that spontaneous fluctuations are undesirable noises, which in principle lead to deterioration of the performance. Considering the significant magnitude of intrinsic dynamics [32,57], however, one could suspect an opposite case where intrinsic dynamics are more beneficial than harmful. Below we review this possibility for representing probabilistic information, maintaining memory, or acquiring new information.

### Statistical inference via synaptic stochasticity

One possible role of intrinsic dynamics is regarding probabilistic computation in the brain. As we all know, life is full of uncertainty: the external world is innately irregular, and available information on the environment is always imperfect and noisy. It is natural, then, to suppose that the brain can efficiently represent and manipulate stochasticity, and many experiments have indeed verified that the brain conducts probabilistic inference at near Bayesian-optimal level [58]. There are several proposals on how probabilistic computation is implemented in the brain [59,60] and the sampling hypothesis, which states that a “snapshot” of a neural circuit is a draw from the encoded distribution [61], is drawing attention among them. While many works focus on the sampling of neural activities (e.g. stochastic firing of neurons [62] or variability of membrane potentials [63–65]), the brain might also perform sampling of circuit configuration via intrinsic dynamics (Figure 2B). Two sampling paradigms differ in timescales (one is from milliseconds to seconds, the other is from hours to days), and the brain might utilize both in a complementary manner. Below we introduce several examples.

In a series of works by Kappel et al. [66,67], they modeled synaptic dynamics as a variant of Langevin sampler [68], which asymptotically represents the posterior distribution. These dynamics allowed synaptic parameters, although changing continuously and stochastically, to remain mostly within a high-dimensional region of high posterior probability (in the setting of unsupervised learning [66]) or large expected reward (in the setting of reinforcement learning [67]), leading to high computational performance in the presence of ongoing synaptic dynamics. The derived synaptic update algorithm reproduced a (reward-modulated) STDP plasticity rule [69,70] as well as showing good generalization ability and robustness to perturbation due to its Langevin dynamics.

Another recent work by Teramae [71] integrated two different timescales of sampling in synapse and neuron. It generalized Boltzmann machine [72] learning with additional weight sampling, and proposed that fluctuations of both neural activity and synaptic weights arose as a result of the Gibbs sampling [73]. The derived algorithm reproduced several non-trivial experimental findings such as the power-law coding of the cortex [74] at the neural activity level and was consistent with the STDP rule at the synaptic level.

### Homeostatic effect of intrinsic dynamics

Recent findings, both experimental and theoretical, have been revealing that intrinsic dynamics could contribute to homeostasis of the neuronal network in two distinct manners, stability of cell assembly and the robust shape of synaptic weight distribution. Although Hebbian plasticity can successfully explain the formation of cell assembly via learning, maintaining them during spontaneous activity has been remaining a challenging problem for conventional learning rules. In a recurrent network, formed assemblies tend to become unstable in additive Hebbian plasticity models due to positive feedback effect [75,76] and degenerative in multiplicative Hebbian plasticity models [77]. Several works have introduced different mechanisms for stabilization [78] such as inhibitory plasticity [75], homeostatic plasticity [79], heterosynaptic plasticity [76], or embedding limit cycles as memory patterns [80], but these models often yield synaptic weight distributions which are far different from experimental observation.

Many experimental works have consistently reported that the distribution of synaptic weights (or the corresponding dendritic spine volumes) have a unimodal, heavy-tailed, and skewed shape (for review, see [81–83]; note that the tail of the distribution is cut off at the upper bounds of spine size (~1μm^3^)). This stereotypical shape seems almost universally preserved, even in the extreme conditions where neuronal networks developed in the absence of synaptic transmission [84,85], spiking activity [29,57], or expression of key molecules for physiological neuronal development [86–88]. It has been suggested that the characteristic shape could play various computational roles, such as efficient information transmission [89,90], enriched repertoires of activity patterns [91], and criticality in the brain [92]. Thus, finding a model of synaptic dynamics which is consistent with this distribution of synaptic weights is necessary for an accurate description of the neural circuit.

A synaptic plasticity model with intrinsic dynamics might be an answer for these two challenges. First, the intrinsic dynamics model explains the robustness of weight distribution in the following way. According to experimental observation [29], the dynamics of synaptic weight are well described with the Langevin equation *dw* = *μ*(*w*)*dt* + *σ*(*w*)*dB_t_* (where *B_t_* represents standard Brownian motion). The drift term *μ*(*w*)*dt* and diffusion term *σ*(*w*)*dB_t_* represent, respectively, extrinsic dynamics (activity-dependent plasticity) and intrinsic dynamics (activity-independent fluctuation). The stable distribution that is computed from the measurement of *μ*(*w*) and *σ*(*w*) reproduced the actual distribution of synaptic weights in both physiological [28,29,57] and pathological conditions [86] (Figure 2C). The analysis of the diffusion process yields that skewness and heavy-tail appear robustly for a wide range of activity-dependent plasticity terms, as long as the magnitude of intrinsic dynamics, *σ*(*w*), is approximately proportional to synaptic weight [9,29].

Several modeling works have also consistently demonstrated that the combination of intrinsic and extrinsic dynamics can lead to stable maintenance of memory [54,55,93]. In particular, a work by Humble et al [93] hinted that intrinsic dynamics are more than noise to be continually corrected by extrinsic dynamics. In their settings where memory is represented as cell assemblies in a recurrent network and synaptic changes are driven by STDP and intrinsic dynamics, intrinsic dynamics alleviate the positive feedback effect of STDP by preventing unnecessary synapses from growing. Further analysis has revealed that an appropriate combination of STDP and intrinsic dynamics could introduce bistability in a group of synaptic weights [9]. The combination created a separation point between large and small synapses to minimize their interconversion, separating strong intra-assembly and weak non-intra-assembly synapses while roughly confining the synaptic weight distribution to a stereotypical shape (Figure 3A). Although the influence of having cell assemblies on the weight distribution looks small in the standard linear plot, it is magnified and form a secondary peak in the semi-logarithmic plot (Figure 3B), which was experimentally detected only recently by the advancement of serial section electron microscopy (ssEM) [42,94]. These findings are reminiscent of older theory on binary synaptic states [95], which exhibited high-performance in ANNs [96], suggesting that the brain might utilize two ‘representative’ synaptic weights. An additional important property of the above model is that individual synapses do not take independent binary states—rather, most intra-assembly synapses take the strong state, and most non-intra-assembly synapses take the weak state.

**Figure 3.**
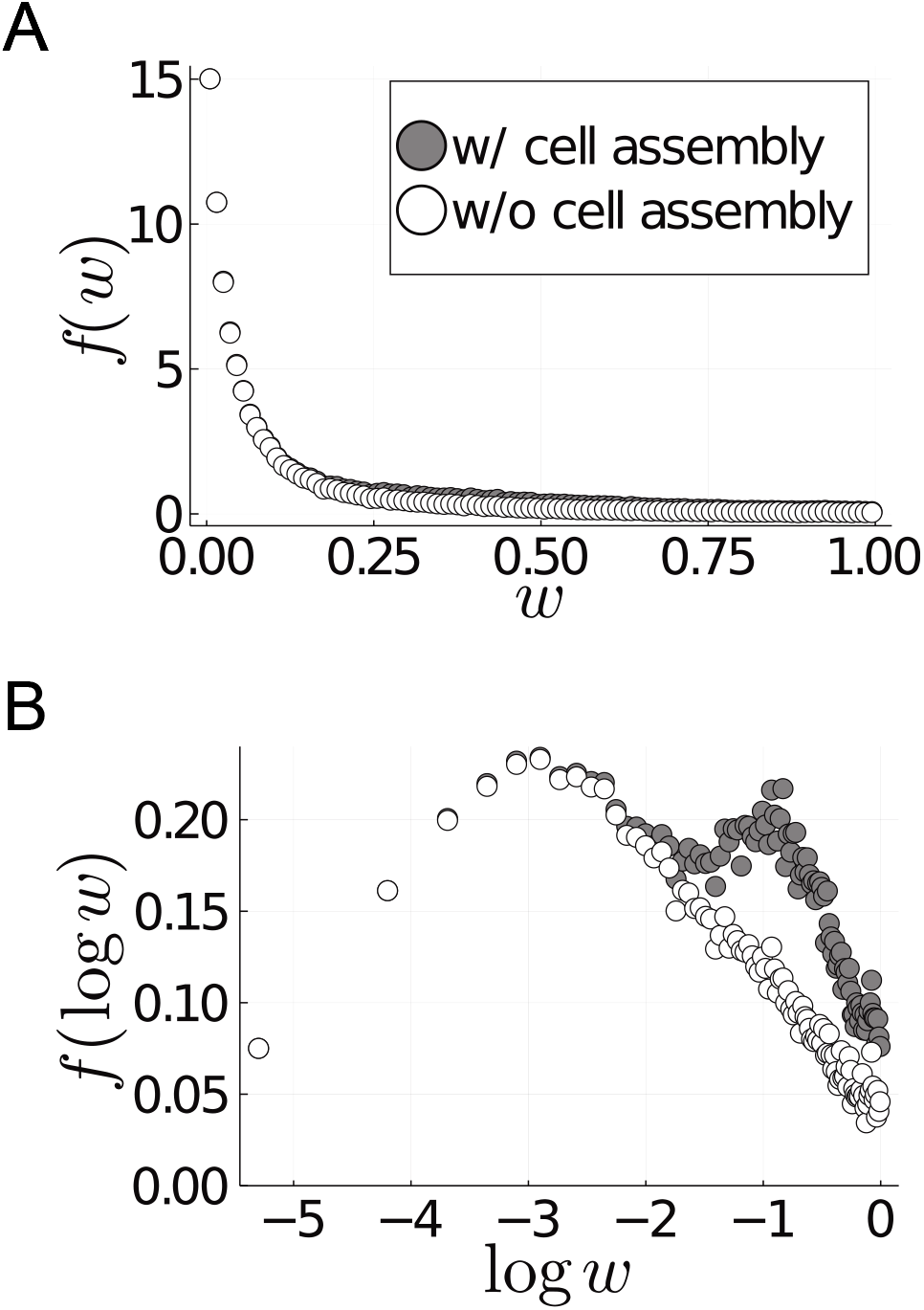
Normal and semi-logarithmic plots of spine sizes and their multi-stability. The data is from Figure 3E of the ref [93], in which synaptic weight *w* is proportional to spine size. A. A standard plot of synaptic weight distribution *f*(*w*). Even after stable memory formation was achieved by the cell assemblies, the model yielded the monotonously decaying stationary distributions for synaptic weights. B. A semi-logarithmic plot of synaptic weight distribution. Following the coordinate transformation, the distribution is given by 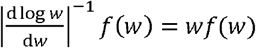. It displayed an additional peak at large synaptic weight if cell assemblies are embedded. This was because the semi-logarithmic plot magnified the nearly invisible difference in (A) for the distributions with and without cell assemblies. Note that the major peak around log *w* =−3.0, which is absent in *f*(*w*), was produced by the coordinate transformation.

### Exploration via Intrinsic dynamics?

Another underexplored potential of intrinsic synaptic dynamics is to enhance exploration. Since the exploration-exploitation trade-off is a key concept in reinforcement learning, engineeringly many algorithms have been proposed to ensure the exploration of the agent. Among them, adding random (and reward- or environment-independent) noise to the policy parameters is simple to implement and has exhibited impressive performance in various settings [97–99]. The advantage of perturbations for enhanced exploration has also been studied in neuronal models, but these studies sought the origin of perturbation for the exterior of the learning network [100], stochasticity of presynaptic vesicle release [101], or intrinsic excitability of neurons [102,103]. While direct evidence remains elusive, synaptic fluctuations might be a novel mechanism to implement trial-and-error exploration of better neuronal networks.

### A link to another dynamics in the brain: representational drift

So far we have mainly discussed brain dynamics at the synaptic level. However, other dynamic features of information processing in the brain are drawing attention as well. In particular, several experimental works have demonstrated that neuronal representation, which refers to the neural activity specifically related to each memory, is continuously drifting over time (for reviews of the relevant literature, see [104,105]). The drift seems ubiquitous across many brain areas including the hippocampus, sensory cortex, and motor cortex, even when the animal shows no apparent change in its behavior. The cause and role of representational drift largely remain unknown. Some modeling works have proposed the framework in which extrinsic synaptic dynamics change neural codes to be consistent with the learning criterion and intrinsic dynamics add isotropic random changes, thereby the representation drifts in the “neutral” dimensions of the learning criterion [67,106]. For example, the network model of [67] was driven by both intrinsic dynamics and the gradient of expected reward. As a result, while the action and rewards were almost identical over trials after learning completion, the neural representation encoding such action was continuously drifting. Experimental evidence to support this idea is not sufficient at present, but the combination of intrinsic and extrinsic synaptic dynamic might be crucial for the brain to be both adaptive and flexible.

## Conclusion

In this work, we reviewed recent developments, based on both experimental and modeling tasks, on the computational roles of intrinsic synaptic dynamics. While conventional rules of neural plasticity attempt to describe synaptic weight changes based on neural activities, here we propose that the widespread presence of activity-independent synaptic fluctuations warrants the investigation of intrinsic dynamics to characterize the changes. In some previous works, it is assumed (although often only implicitly) that intrinsic dynamics do not compromise network performance. Different insights from neuroscience and machine learning, however, raise the possibility that brain dynamics not only are constantly affected by intrinsic dynamics but could benefit from their presence. Future theoretical and experimental studies might give us an understanding of why our brain evolutionarily has chosen to develop neural networks based on intrinsically dynamical components.

## Conflict of interest statement

The authors declare no conflicts of interest.

## References and recommended reading

Papers of particular interest, published within the period of review, have been highlighted as:

* of special interest
** of outstanding interest

## Acknowledgements

We thank M. Baltieri for proofreading the manuscript and K. Ichikawa for helpful discussions. The study was supported by RIKEN Center for Brain Science, Brain/MINDS from AMED under Grant No. JP21dm0207001 (T.T.), KAKENHI Grant-in-Aid JP18H05432 (T.T.) and 20H05685 (H.K.) from JSPS, RIKEN Junior Research Associate Program (K.Y.), Masason Foundation (K.Y.), WPI from MEXT (H.K.), CREST JPMJCR1652 (H.K.) from JST, and SRPBS JP20dm0107120 (H.K.) from AMED.

## (annotations for selected literature)

[29] (Yasumatsu et al., J Neurosci 2008)

(*) The first report of intrinsic synaptic dynamics. The authors observed, using chronic two-photon imaging, that dendritic spine volumes fluctuated even after the total pharmacological blockade of neural activities in hippocampal slice cultures, and the fluctuations quantitatively account for the statistics of spine volumes using stochastic differential equations.

[31] (Nagaoka et al., Sci Rep 2016)

(*) The first study to report that intrinsic dynamics is also present in vivo, using chronic two-photon imaging with cortical activity silenced only locally. They also reported that enhanced synaptic turnover rate in Fmr1 KO model mice primarily reflected an elevated level of intrinsic dynamics.

[38] (Mongillo et al., Nat Neurosci 2018)

(**) The authors showed in the balanced network model of a cortical circuit that rearranging inhibitory connectivity more significantly altered network activity patterns than excitatory connections in the condition that the time-averaged firing rate is higher and more diverse in inhibitory neurons. In applying this argument to memory maintenance and recall they assumed anti-Hebbian plasticity in inhibitory synapses.

[42] (Dorkenwald et al., bioRxiv 2019)

(*) The large connectivity map between layer 2/3 pyramidal cells in the mouse primary visual cortex was presented using the ssEM technique. They showed synapse sizes could be modeled by the sum of a binary variable and an analog variable drawn from a log-normal distribution and suggested that the former result from activity-dependent plasticity rule while the latter reflect intrinsic dynamics. [52] (Dubreil et al., bioRxiv 2020)

(*) In order to unify two competing views on neural computation, functional cell classes and collective dynamics, the authors developed a novel class of interpretable recurrent networks. After the network was trained for each task, the minimal dimension of intrinsic collective dynamics and the number of neural sub-populations were estimated. These extracted structures were found essential for task performance in that both (1) resampling connectivity from the estimated sub-populations and (2) reducing to a simple dynamical system consisting of minimal intrinsic dimensions and sub-populations led to comparable performance to the original network. Analysis of the reduced model has revealed that the sub-population structure extends the range of possible dynamics for the internal variables, and in particular, allows for flexible modification of input-output associations.

[54] (Fauth and van Rossum, eLife 2019)

(*) They showed that cell assembly is stably maintained in the presence of random synaptic turnover when spontaneous reactivation reinforces memory during resting periods. This model assumes that intrinsic synaptic dynamics change connectivity but not synaptic weight.

[55] (Acker et al. J Neurophysiol 2019)

(*) They showed that continual correction by Hebbian plasticity is sufficient to preserve neural representation in the presence of random spine turnover in the model of (1) the grid-cell-to-place-cell transformation in CA1 and (2) the center-surround-to-simple-cell transformation in V1.

[64] (Echeveste et al., Nat Neurosci 2020)

(*) In the article, the authors trained a canonical balanced neuronal network model of the sensory cortex for a sampling-based inference and found the optimized network exhibited various cortical-like dynamical features, such as gamma-oscillations, stimulus-modulated noise variability, transient overshoots at stimulus onset.

[65] (Aitchison et al, Nat Neurosci 2021)

(*) The authors propose a novel synaptic plasticity rule in which both synaptic weights and their uncertainty are updated via learning. The proposed normative rule shows a larger learning rate when the uncertainty about synaptic weight is larger; in other words, the less confidence a synapse has in its target weight, the larger influence new information has on its update. They further suggest that this uncertainty is represented as the variability in the postsynaptic potential size.

[67] (Kappel et al., eNeuro 2018)

(**) The authors proposed a framework to regard synaptic fluctuations as sampling dynamics. Via a combination of reward-modulated plasticity and intrinsic dynamics, the network could move mostly within the configuration region of high expected reward under ongoing synaptic changes.

[71] (Teramae. bioRxiv 2020)

(**) The author hypothesized that the brain generated synaptic and neural states that were consistent with the external world. More precisely, they formulated that both the firing of neurons and formation of synapses were the result of Gibbs sampling conditioned on the external environment and the status of other neurons and synapses. This perspective could be regarded as the extension of Boltzmann machine learning with additional synaptic sampling. This framework predicted the existence of retrograde modulation by postsynaptic neurons, as well as consistently explained various experimental findings in the brain.

[80] (Susman et al., Nat Commun 2019)

(*) Hinted by general theory on system stability, this study suggests that the anti-symmetric component of the connectivity matrix could be more stable than the symmetric component when activity-dependent homeostatic plasticity predominantly affects the symmetric component. They further proposed an algorithm to implement memory encoding and decoding in this anti-symmetric domain by extending the Hopfield network model.

[86] (Ishii et al., eNeuro 2018)

(*) The authors recorded spine dynamics of Fmr1 KO mice, which are one of the animal models of ASD, in the visual cortex using in vivo chronic two-photon imaging. They reported that Fmr1 KO mice showed a higher rate of spine fluctuations for the first time in vivo, which led to the predominance of smaller spines in Fmr1 KO mice.

[92] (Kusmierz et al., Phys Rev Lett 2020)

(*) A novel proposal for the computational role of heavy-tailed synaptic distribution. The authors studied a random network model in which synaptic weights are drawn independently from the identical Cauchy distribution. The network showed a continuous phase transition from a quiescent state to a chaotic state. Neuronal avalanches (i.e., bursts of activity with power-law distributions of sizes and lifetimes) were reproduced at the critical point (i.e., the edge of chaos). By contrast, the Gaussian counterpart showed a discontinuous jump to chaos under the assumption that neurons have a firing threshold and receive many synaptic inputs, thus could not be posed robustly at the edge of chaos.

[93] (Humble et al. Front Comput Neurosci 2019)

(**) They showed combinations of Hebbian plasticity and intrinsic dynamics stabilized cell assembly while maintaining the physiological heavy-tailed distribution of synaptic weight. This stable maintenance of cell assembly was impaired by strong intrinsic dynamics observed in some ASD models.

[98] (Fortunato et al., ArXiv 2019)

(**) The authors introduced NoisyNet, a deep reinforcement learning agent with probabilistic noises added on its weights aiming at efficient exploration. Although straightforward to implement, this agent showed substantially better performance on a wide range of Atari games than conventional heuristic methods for exploration.

[106](Kalle-Kossio et al., bioRxiv 2020)

(**) In this modeling work, the authors suggested that representational drift was driven by spontaneous synaptic remodeling which caused a transition of neurons between cell assemblies, while consistency of inputs – assembly – outputs relationship was maintained via Hebbian plasticity without requiring any additional teaching signals or behavioral feedback. They also reported that probabilistic firing of neurons contributed to drift as well, suggesting the importance of integrating the stochasticity of different levels.

## Notes

### Competing Interest Statement

The authors have declared no competing interest.

